# Human NK cells engage cell-intrinsic non-canonical inflammasome function

**DOI:** 10.1101/2025.10.20.682800

**Authors:** Nathan G.F. Leborgne, Jadie Acklam, Alexander J. Hogg, Zarema Albakova, Karen G. Hogg, Kim Samirah Robinson, Grant Calder, Arnel Villamin, Callum Robson, Freya Pealing, Matthew A. Care, Monika Gonka, James M. Fox, James Iremonger, Frances E. Pearson, Max E. Noble, Alison M. Layton, James P. Hewitson, David G. Kent, William Grey, Jillian L. Barlow, Dave Boucher

## Abstract

Inflammasome pathways are critical for detecting pathogens and initiating inflammatory responses, yet how individual cell types configure these signalling platforms remains poorly understood. Here, we demonstrate that human Natural Killer cells possess distinct inflammasome machinery, lacking canonical components and innate sensors (NLRP3, NLRC4, TLR4) but harbour a functional non-canonical inflammasome (NCI). Using *Salmonella* enterica Typhimurium infection of primary human NK cells, we show that cytosolic bacterial access triggers caspase-4-dependent pyroptosis and IL-18 secretion, without canonical inflammasome activation or IL-1β production. Strikingly, murine NK cells express inflammasome transcripts but lack functional caspase-11 protein and do not undergo pyroptosis upon infection, revealing species-specific post-transcriptional regulation. We further identify IL-12, but not IFNγ, as a priming signal for NCI components in human NK cells, a functional distinction from macrophages and epithelial cells. Together, these findings reveal that human NK cells have evolved a specialised inflammasome with unique regulatory inputs, establishing these cells as autonomous sensors of intracellular Gram-negative infection.

## Introduction

Innate immune responses are controlled by cellular pathways that coordinate immunity to rapidly eliminate pathogens and restore homeostasis (1). Inflammasomes are critical signalling platforms controlling responses to sterile and infectious agents in immune and epithelial cells (2,3). The inflammasome pathway senses triggers using pattern-recognition receptors (PRRs), which include members of the Nod-like Receptor family (Nod-like receptor protein-1, NLRP1, and −3, NLRP3) and the NLR family CARD domain-containing protein 4 (NLRC4) (2). Unlike the canonical inflammasome, which activates caspase-1 upon detection of bacterial ligands (*e.g.,* flagellin), and self-derived danger signals (*e.g.,* ATP, nucleic acids), the non-canonical inflammasome (NCI) activates caspase-4 and caspase-5 (humans) and caspase-11 (mice) upon recognition of cytosolic bacterial lipopolysaccharides (LPS) (4–6) or from intracellular delivery of outer membrane vesicles (7). In non-phagocytic cells, such as epithelial cells, LPS is presented to caspase-4 through interferon-inducible proteins called guanylate-binding proteins (GBPs) (8–12). GBP1 is a direct LPS sensor and is involved in NCI activation in human cells upon *Salmonella* infection (9,12). NCI signalling leads to the maturation and secretion of pro-inflammatory cytokines (13) and to pyroptosis, a form of pro-inflammatory cell death (14). Pyroptosis is driven by the caspase-mediated cleavage of the pore-forming protein Gasdermin D (GSDMD) (14–16), which activates a Ninjurin-1 (NINJ1)-dependent plasma membrane rupture (17), and is critical to control infections, limiting their spread and mobilising further immune cells for pathogen clearance (18). Pyroptosis mediates the release of antigens, which support adaptive responses (19). Additionally, GSDMD pores mediate potassium influx, which activates the NLRP3 canonical inflammasome downstream of caspase-4/11 (20–22). Inflammasome activation in macrophages and epithelial cells induces the maturation and secretion of interleukin-18 (IL-18) and IL-1β, which are released through the GSDMD pores (23,24). These cytokines generate antimicrobial responses in myeloid, lymphoid, and epithelial cells (25).

Discovered in the 1970s, NK cells are part of the now-expanded family of innate lymphoid cells (ILCs), including ILC1s, ILC2s, ILC3s, and lymphoid tissue-inducer cells. They play key roles in viral and tumour responses and are increasingly studied *ex vivo* as part of novel immunotherapy against cancer (26,27). Functionally, these cells are broadly divided into cytotoxic NK cells (release of perforin, granzyme and defined as CD56^dim^CD16^+^) and immunoregulatory NK cells (IL-12- and IL-18-dependent release of proinflammatory cytokines, IFNγ, CCL3, GM-CSF and defined as CD56^high^CD16^low^) (28,29). Progress has been made in recent years on the contribution of NK cells to early responses during microbial infection. Several studies have shown that NK cells indirectly respond to bacterial infection via IL-12 and IL-18 secreted by accessory cells such as macrophages and dendritic cells, leading to IFN-γ production. (30–32). Human NK cells can also directly recognise bacterial PAMPs, such as outer membrane protein A (OmpA), flagellin, and bacillus Calmette-Guérin (BCG), via TLR2, TLR5, or NKp44, respectively, triggering NK proliferation, IFN-γ production, and α-defensin release without requiring accessory cells (33,34). However, our knowledge of direct bacterial recognition by NK cells remains limited.

Inflammasomes influence adaptive responses in NK, B, and T cells through cytokine-driven and pyroptosis-mediated antigen release from myeloid and epithelial cells (19). As such, inflammasomes are an important part of strong immune responses to vaccines and long-term defence against pathogens (35,36). Recently, cell-intrinsic inflammasome-dependent pyroptosis was demonstrated in naive T cells (37,38). Moreover, it is now established that inflammasome derived-IL-18 can activate NK cells and promote the expression of interferon-γ (IFNγ) and cytotoxic effectors (39). Thus, NK cells are currently viewed as a downstream effector in the inflammasome pathway.

To date, inflammasome functions have mainly been studied in myeloid and epithelial cells where Gram-negative bacteria, such as *Salmonella*, are detected through exposure to a range of Pathogen-Associated Molecular Patterns (PAMPs), including type III secretion system components, flagellin and LPS (40,41). Whether NK cells possess effector and cell-intrinsic inflammasome activity has not been investigated.

Here, we investigate how inflammasomes shape NK cell behaviours during infection with the Gram-negative bacteria *Salmonella enterica* Typhimurium. Using primary human NK cells isolated from blood, we identified a unique inflammasome signature that drives NCI activation but not canonical inflammasome activation. This signature was further enhanced upon IL-12 priming and was absent in mouse NK cells. Our findings highlight the unique regulation and roles of the inflammasome in human NK cells and describe direct intracellular sensing functions during gram-negative infections.

## Results

### Human NK cells possess a functional non-canonical inflammasome

We took advantage of a substantial dataset of healthy human blood NK cells to profile the expression of inflammasome pathway components in these cells where gene and protein expression data in human blood NK cells from 13 healthy donors (36,270 high-quality filtered cells) were grouped functionally as, NK1 (CD56^dim^ CD16^high^CD3^-^CD127^-^CD57^-^CD2^-^, further divided as subtype A, B, and C), NK2 (CD56^high^CD16^-^CD3^-^CD127^+^CD57^-^ CD2^+^), NK3 (CD56^dim^CD16^high^CD3^-^CD127^-^CD57^+^CD2^-^), and NKint (42). As expected, *GZMB, GZMA,* and *PRF1 showed* higher mean expression in NK1 and NK3 cells (**Figure 1A**). Strikingly, canonical inflammasome components (*NLRP3*, *NLRC4*), the pro-inflammatory cytokine IL-1β, and the receptor detecting extracellular LPS (*TLR4*), were not expressed (40,43,44). *CASP1* and *GSDMD* were detected in NK1 and NK3, but not in NK2 subtype. However, we were able to observe increased mean expression of NCI components, including *CASP4*, *GBP1*, and *GBP2*, in NK1 and NK3 cells, compared to NK2 cells. Other inflammasome-related genes were further restricted, with *GBP4* expressed only in NK1B and NK2, and *PYCARD* in NK1B and NK1C (**Figure 1A**). To exclude the possibility that this was blood-specific, we examined gene expression in human foetal thymic (tissue-resident) NK cells and confirmed that *NLRP3*, *NLRC4*, and *TLR4* were not expressed. We identified a *CASP4^+^GBP2^+^*NK cell population, in which *CASP1, GSDMD*, and *PRF1* were upregulated in cycling cells, as compared to non-cycling cells (**Supp. Figure 1A**). This suggests the ability of specific NK cell populations to respond to intracellular Gram-negative infections in different tissues.

**Figure 1.**
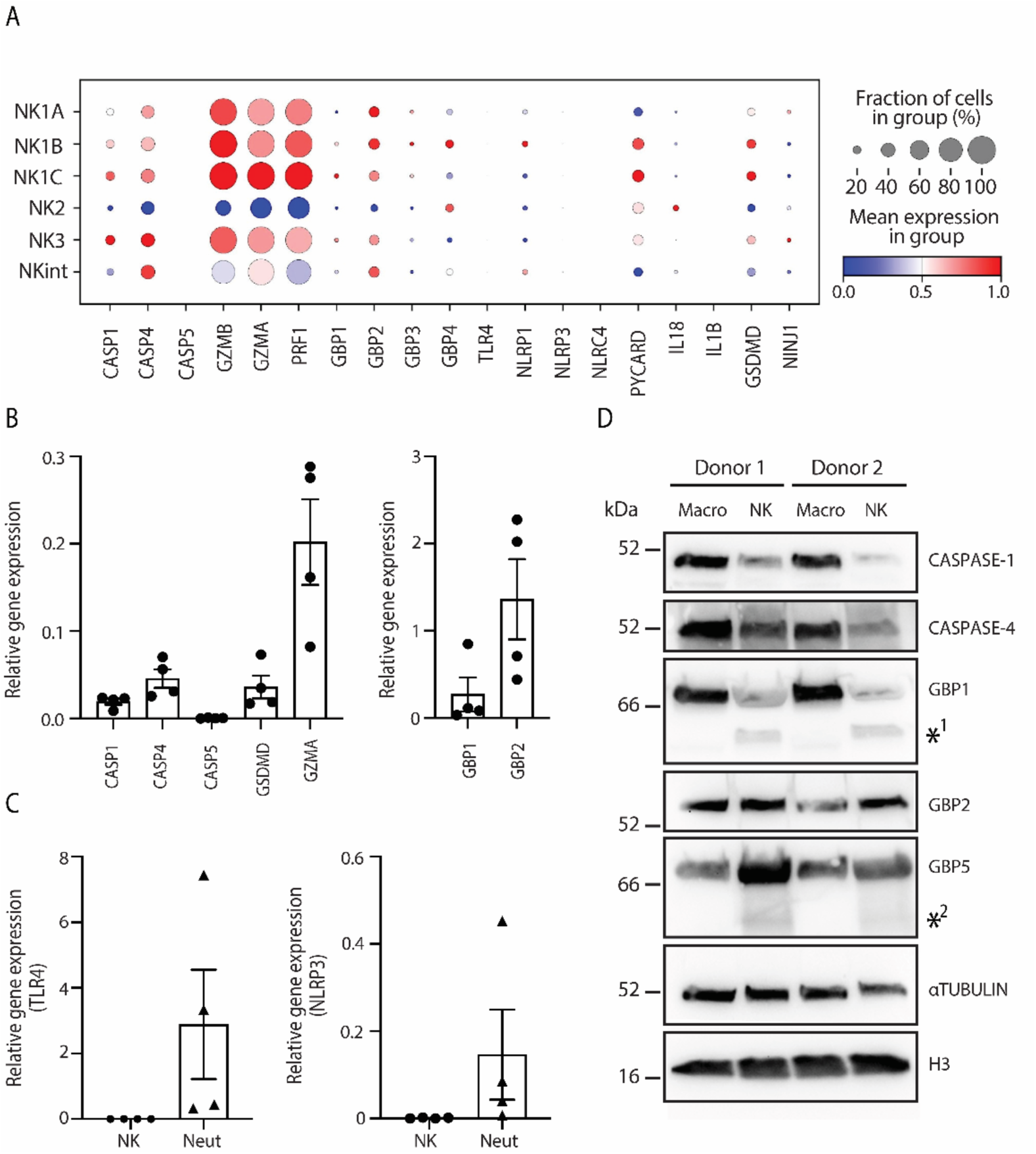
Human blood NK cells express components of the non-canonical inflammasome. (**A**) scRNA-seq data of inflammasome components in subsets of human peripheral blood NK cells showing the relative abundance (standard-scaled) and percentage of cells expressing inflammasome components, (n=19). (**B**) Quantitative PCR analysis of non-canonical inflammasome components in unexpanded human peripheral blood NK cells, n = 4 donors (**C**) Quantitative PCR analysis of *TLR4* and *NLRP3* in unexpanded peripheral blood NK cells and neutrophils from four matched donors. (**D**) Western blot analysis of expanded human peripheral blood NK cells and macrophages from 2 matched donors compared to loading controls of αTubulin and H3. *^1^ and *^2^ indicate NK cell-specific protein isoforms of GBP1 and GBP5, respectively. Western blot is representative of 4 blood donors.

To confirm the findings from these datasets, we next isolated human peripheral blood NK cells and neutrophils, paired from the same donor and performed quantitative PCR for inflammasome-related genes. Alongside GZMA expression, NK cells also showed low expression of *CASP1*, *CASP4*, and *GSDMD*, with greater expression of *GBP2* (**Figure 1B**). This was compared to expression of these genes in neutrophils, but whilst neutrophils showed gene expression for *TLR4* and *NLRP3,* NK cells did not (**Figure 1C and Supp. Figure 1B**). Analysis of human blood NK cells, using macrophages as a comparison, by western blot also indicated lower protein expression of caspase-1, caspase-4, and GBP1 in NK cells, and similar levels of GBP2 and GBP5. We identified NK-specific proteoforms of GBP1 and GBP5 (**Figure 1D**), also previously reported in HUVEC cells (45). Overall, these data reveal that human NK cells operate a minimalist inflammasome system, lacking the canonical machinery (NLRP3, NLRC4, TLR4) that defines macrophage responses to *Salmonella*. Instead, this suggests they rely exclusively on the NCI pathway for bacterial detection.

To test the activation of the NCI in the context of infection, we next expanded negatively-selected human peripheral blood NK cells *in vitro* for 14 days, resulting in a highly enriched population containing all NK cell subpopulations, as compared to NK cells directly isolated from fresh blood (**Supp. Figure 2**). We infected the NK cells with *Salmonella enterica* Typhimurium, strain SL1344 and LT2, which infect cell types such as macrophages and epithelial cells, activating canonical (NLRP3, NLRC4; caspase-1) and non-canonical (GBPs; caspase-4,-5) inflammasomes. Infection resulted in a statistically significant increase in lytic cell death, as measured by LDH release (**Figure 2A**), compared to uninfected cells. This cell death was decreased with the addition of the caspase-1/-4 inhibitor, VX-765 (46) (**Figure 2A**) and upon infection with *SalΔorgA,* a *Salmonella* mutant lacking the functional SPI1 injectisome required for the invasion of non-phagocytic cells (47) (**Figure 2B**). Indeed, *Salmonella* enter the cytosol (9,10) by using a subset of effectors (48), including the needle injectisome protein, OrgA, where its deletion strongly reduces *Salmonella* invasion of non-phagocytic cells (40). Thus, our data not only suggest a caspase-1/-4-dependent cell death mechanism, but also that NK cell lytic death requires intracellular *Salmonella* invasion.

**Figure 2.**
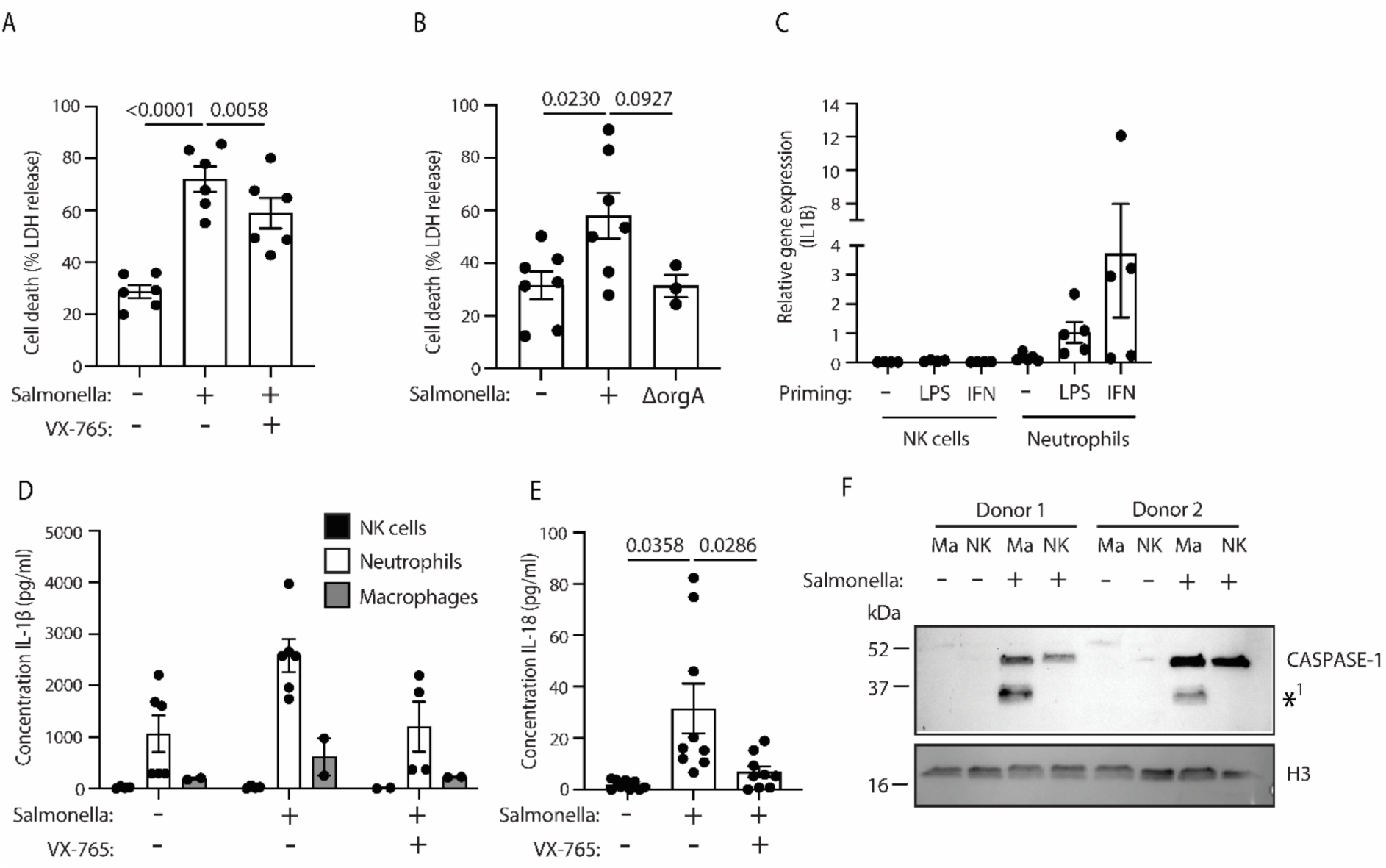
*Salmonella* Typhimurium infection induces cell death and IL-18 production in human blood NK cells. Expanded human peripheral blood NK cells were infected with *S.* Typhimurium or *S.* Typhimurium*ΔorgA*, and with or without VX-765. At 5 hrs post-infection, cells supernatants were assessed by (**A** and **B**) LDH assay for cell death, and (**D** and **E**) ELISA for IL-1beta and IL-18 production, n = 6 - 9 donors across 2 - 3 independent experiments. (**C**) Quantitative PCR analysis of *IL1B* following priming with LPS (for 3 hrs) and IFNgamma (for 16 hrs). **(F)** At 5 hrs post-infection, caspase-1 cleavage was characterised by western blot. *^1^ indicate a p33 cleavage product of active caspase-1. In (**C**), (**D**) and (**F**) NK cells were compared to neutrophils and macrophages from matching donors. Data in (A, B and E) were analysed by one-way ANOVA with Dunnett’s multiple-comparison test. p values are displayed and p<0.05 was considered statistically significant.

Central to the activation of the NCI is the maturation and secretion of pro-inflammatory cytokines, such as IL-1ꞵ and IL-18. Our data confirm that priming with LPS or IFNγ, or infection with *S.* Typhimurium induces IL-1ꞵ gene expression (**Figure 2C**) and its secretion (**Figure 2D**) in human neutrophils and macrophages. However, IL-1ꞵ was not detectable in NK cells under the same conditions in matched donors. Strikingly, *S.* Typhimurium infection of NK cells induced a statistically significant increase in IL-18 production, which was caspase-1/-4-dependent (**Figure 2E**).

As *Salmonella* has been shown to activate multiple inflammasomes during infection, we investigated the pathways involved in such a response. Interestingly, we could not detect the cleaved caspase-1 fragments (p20/p10) upon *Salmonella* infection (**Figure 2F**). Additionally, Nigericin treatment induced an NLRP3-independent cell death in NK cells, as shown with the addition of its inhibitor, MCC950 (49) (**Supp. Figure 3A**). NLRP3-dependent caspase-1 cleavage was also not observed in NK cells (**Supp. Figure 3B**). Moreover, Val-boroPro (VbP, NLRP1 inflammasome activator, (50)) did not lead to GSDMD cleavage in NK cells but was observed in a keratinocyte cell line (N/TERT) (**Supp. Figure 3C - D**)

Our data suggest a distinct mechanism of *Salmonella* sensing in NK cells that does not involve canonical inflammasome signalling, as evidenced by the undetectable gene expression level of NLRC4 and NLRP3 and the absence of response to NLRP3 and NLRP1 triggers. Instead, NK cells utilise a pathway that leads to inflammatory caspase-dependent pyroptosis, and IL-18 secretion, but not IL-1ꞵ. In addition, the absence of *TLR4* expression in NK cells, which has been shown to be important for priming the canonical inflammasome pathway, may explain NK cell non responsiveness to NLRP3 triggers and the absence of IL-1β. This differs from the classical activation of these pathways in macrophages where NLRC4 and NLRP3 are major contributors to the innate detection of *Salmonella* (*43,51,52*). The inflammatory caspase-dependent secretion of IL-18 is consistent with previous literature showing its cleavage by caspase-4 (9,10,53–55) and suggests that NK cells use an autocrine mechanism to potentiate their activation during bacterial infection.

### *Salmonella* Typhimurium accesses the cytosol in human NK cells

Gram-negative bacteria can activate the NCI through a range of mechanisms, including direct invasion of the host cell cytoplasm or secretion of outer-membrane vesicles (7). Upon infection of epithelial and myeloid cells, *Salmonella* localises and replicates in specialised vacuoles, where it is protected from immune detection. However, as established in gut epithelial cells, *Salmonella* can escape these vacuoles, allowing cytosolic hyper-replication but also its detection by cytoplasmic sensors (56). Therefore, we next sought to identify whether *Salmonella* invade NK cells by infecting them with *S.* Typhimurium LT2-TdTomato. Using flow cytometry, we showed that ∼40% of NK cells were TdTomato^+^ (**Figure 3A**), suggesting an efficient invasion of NK cells. Moreover, fluorescence microscopy on NK cells indicates the presence of Tdtomato^+^ *S.* Typhimurium in both CD56^+^ vacuole-like structures and in the cytosol (**Figure 3B**). We observed a range of NK cell-*Salmonella* interactions as shown in **Supp. Figure 4A**. However, we could not fully differentiate between cytosolic and vacuole-associated *Salmonella*. To confirm that *Salmonella* could access the cytosol, we used *Salmonella* expressing a glucose-6-phosphate-dependent cytosolic eGFP-reporter (9). As glucose-6-phosphate is only accessible in the cytosol, only cytosolic *Salmonella* will express the fluorescent GFP reporter. At 5hrs post-infection, a significant percentage of NK cells (10-15%), were eGFP^+^ (**Figure 3C**), similar to levels in epithelial cells (9), suggesting that *Salmonella* can invade the cytosol of a proportion of NK cells. This correlates with NCI activation, as shown by western blot for the active caspase-4 p32 cleavage product (13) and the reduction in full-length GSDMD in NK cells in response to *S.* Typhimurium (**Figure 3D**). In comparison, infection with a Gram-positive bacteria, *Staphylococcus* aureus, which is known to activate the canonical inflammasome (57), did not induce caspase-4 cleavage or reduce full-length GSDMD protein.

**Figure 3.**
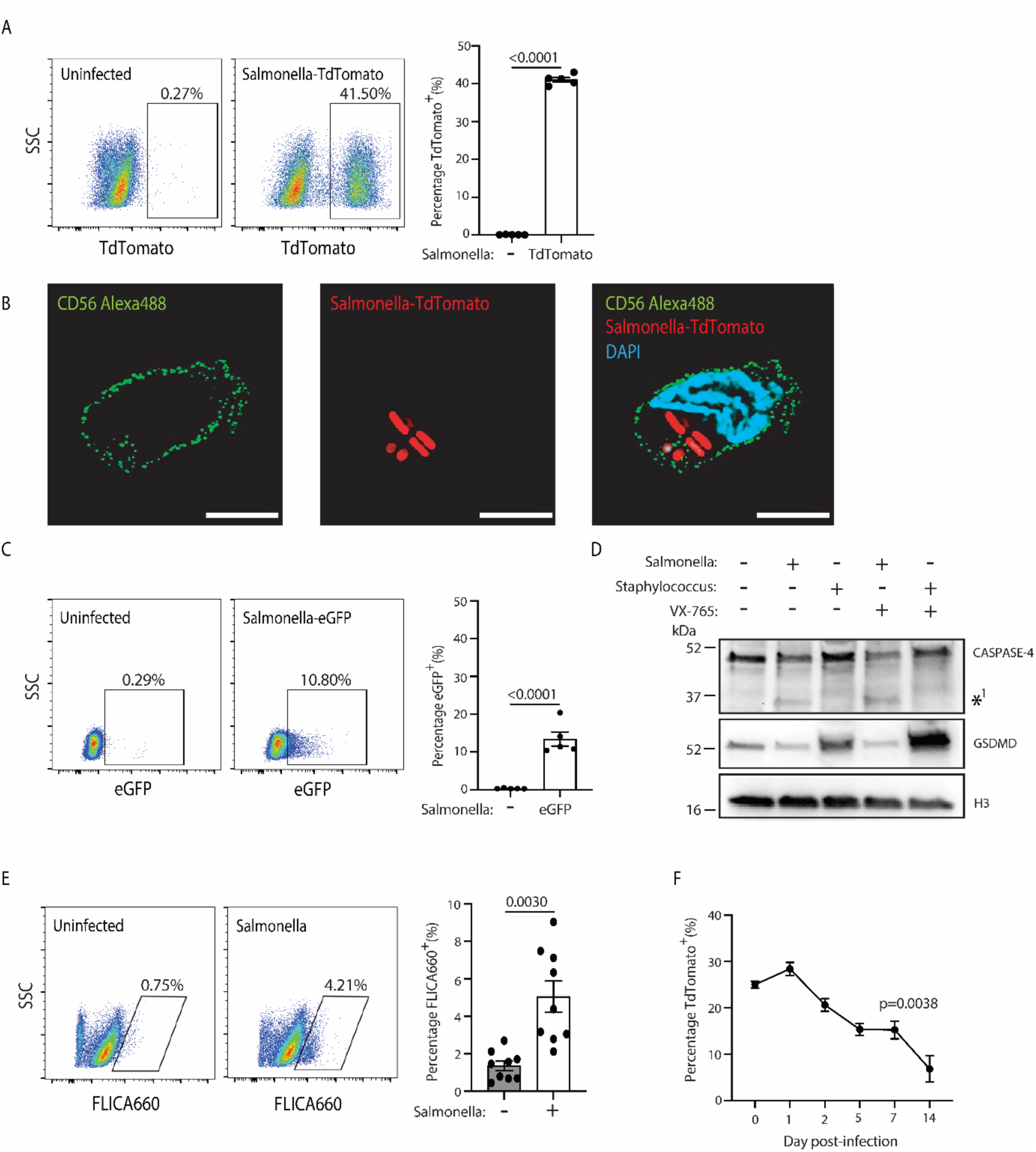
*Salmonella* Typhimurium accesses the cytosol and persists in human blood NK cells. Expanded human peripheral blood NK cells were infected with *S.* Typhimurium-TdTomato. (**A**) At 5 hrs post-infection NK cells were subjected to analysis by flow cytometry or (**B**), stained on fibronectin coated slides with anti-CD56 and DAPI and visualised by fluorescence microscopy (white bar indicates 5 µm). (**C**) Alternatively, these NK cells were infected with *S.* Typhimurium, stably expressing a cytosolic-induced eGFP reporter and analysed by flow cytometry. (**D**) Representative western blot analysis of expanded human peripheral blood NK cells infected with either *S.* Typhimurium or *Staphylococcus* aureus and with or without VX-765 for 5 hrs compared to loading controls of H3. *^1^ indicates a p32 cleavage product of active caspase-4 (**E**) Caspase-1 and −4 activation was measured by flow cytometry using the inflammatory caspase probe FLICA660. (**F**) Percentage of *S.* Typhimurium*-*TdTomato in NK cells was measured at days 1, 2, 5, 7, and 14 post-infection. Data from 2 independent experiments, n = 4 - 6 donors. Data in (A, C and E) were analysed by paired t tests. Data in (F) were analysed by one-way ANOVA with Dunnett’s multiple-comparison test. p values are displayed and p<0.05 was considered statistically significant.

To confirm inflammatory caspase activation, we used the inflammatory caspase activity-based probe FLICA660 and showed that ∼5% of total NK cells were FLICA^+^ upon *Salmonella* infection, compared to ∼1% in uninfected cells, validating caspase-4 activation (**Figure 3E**). In alignment with our previous microscopy and CellView^TM^ analysis indicates a range of activation and infection states, with the existence of FLICA^+^*Salmonella-*TdTomato^-^,FLICA^-^*Salmonella-*TdTomato^+^,and FLICA^+^*Salmonella-*TdTomato^+^ NK cell populations (**Supp. Figure 4C**). To test the potential of NK cells to act as an infectious reservoir for *Salmonella,* we tracked *S.* Typhimurium LT2-TdTomato infection over 14 days. After 5 hrs and 24 hrs, initial infection persists in a significant subset of NK cells, ∼25 - 32%. Between day 1 and day 14 post-infection, this proportion decreases from ∼30-20% to 10-5% (**Figure 3F**), indicating a small NK population in which *Salmonella* persists. Thus, our study suggests heterogeneity in infective states in NK cells, with persistence, potentially in the vacuole, and access to cytosolic niches. Further study will be required to determine whether these differences in persistence and access are associated with specific NK cell subsets. Overall, these mechanisms converge on *Salmonella’s* ability to create an intracellular niche in NK cells.

### Mouse NK cells do not activate the non-canonical inflammasome

We next asked whether murine NK cells displayed similar features, using published RNA-seq datasets (58), where gene expression changes were measured in innate-like lymphoid populations, as compared to macrophages, following LPS priming. Our analyses showed that, in contrast to humans, mouse NK cells express *Tlr4*. Other non-canonical inflammasome-associated genes, such as *Gbp2*, *Casp4*, and *Gsdmd*, were expressed, similarly to human NK cells and other mouse innate lymphoid populations (**Supp. Figure 5A**) (59). These results were supported by qPCR analysis of murine wild-type splenic NK cells (**Supp. Figure 5B**). In naive NK cells, we detected transcripts for *Casp1*, *Casp4*, *Gbp2*, *Gsdmd*, and *Ninj1*, which was comparable to expression seen in macrophages. Macrophages were additionally found to express transcripts for *Il18*, *Il1b*, and *Il1a* (also upregulated with LPS priming), but these transcripts were absent in NK cells (**Supp. Figure 5C**). In contrast to human NK cells, *Salmonella* infection did not induce cell death in murine NK cells (**Supp. Figure 5D**). Furthermore, we did not detect protein expression of full-length caspase-11 (the murine orthologue of human caspase-4) in NK cells compared with matched neutrophils (**Supp. Figure 5F**). Unlike human NK cells, which express inflammasome components at the gene level, murine NK cells do not express a functional caspase-11 and do not activate the NCI in response to *Salmonella* infection.

In contrast, human NK cells do not express TLR4 but instead use the non-canonical inflammasome to detect intracellular bacterial infection. These data highlight a critical difference between human and murine inflammasomes at the cellular level.

### The non-canonical inflammasome in human NK cells is primed by IL-12

IFNγ priming in non-phagocytic and phagocytic cells has previously been identified to impact NCI activation (9–11). To investigate the effect of IFNγ on NK cells, we primed resting NK cells *in vitro* for 16 hrs with recombinant human IFNγ or with the NK-activating cytokine IL-12. This cytokine is also important for the human response to *Salmonella,* as individuals with a deficiency in IL-12 signalling are especially sensitive to *Salmonella* infection (60–63).

Surprisingly, IFNγ did not induce expression of any NCI-related genes (*CASP4*, *CASP1*, *GBP1*, *GBP2*, *GBP5*, *GSDMD*, and *NINJ1*), as compared to unstimulated cells (**Figure 4A - C**). In contrast, IL-12 stimulation resulted in statistically significant increases in *GBP1* and *GSDMD*. Although this did not increase NK cell death, as measured by LDH assay at 5 hrs post-infection (**Figure 4D**), immunoblotting showed increased cleavage of caspase-4 and GSDMD (**Figure 4E**). Ninj1 levels remained unchanged. IL-12 priming significantly increased IFNγ and CCL3 secretion by NK cells, but did not further enhance this response upon *S.* Typhimurium infection (**Figure 4F and G**). Importantly, the *Salmonella*-dependent IL-18 secretion previously observed (**Figure 2E**) was increased upon IL-12, but not IFNγ priming, indicating a potentiation of NCI activity in human NK cells (**Figure 4H**).

**Figure 4.**
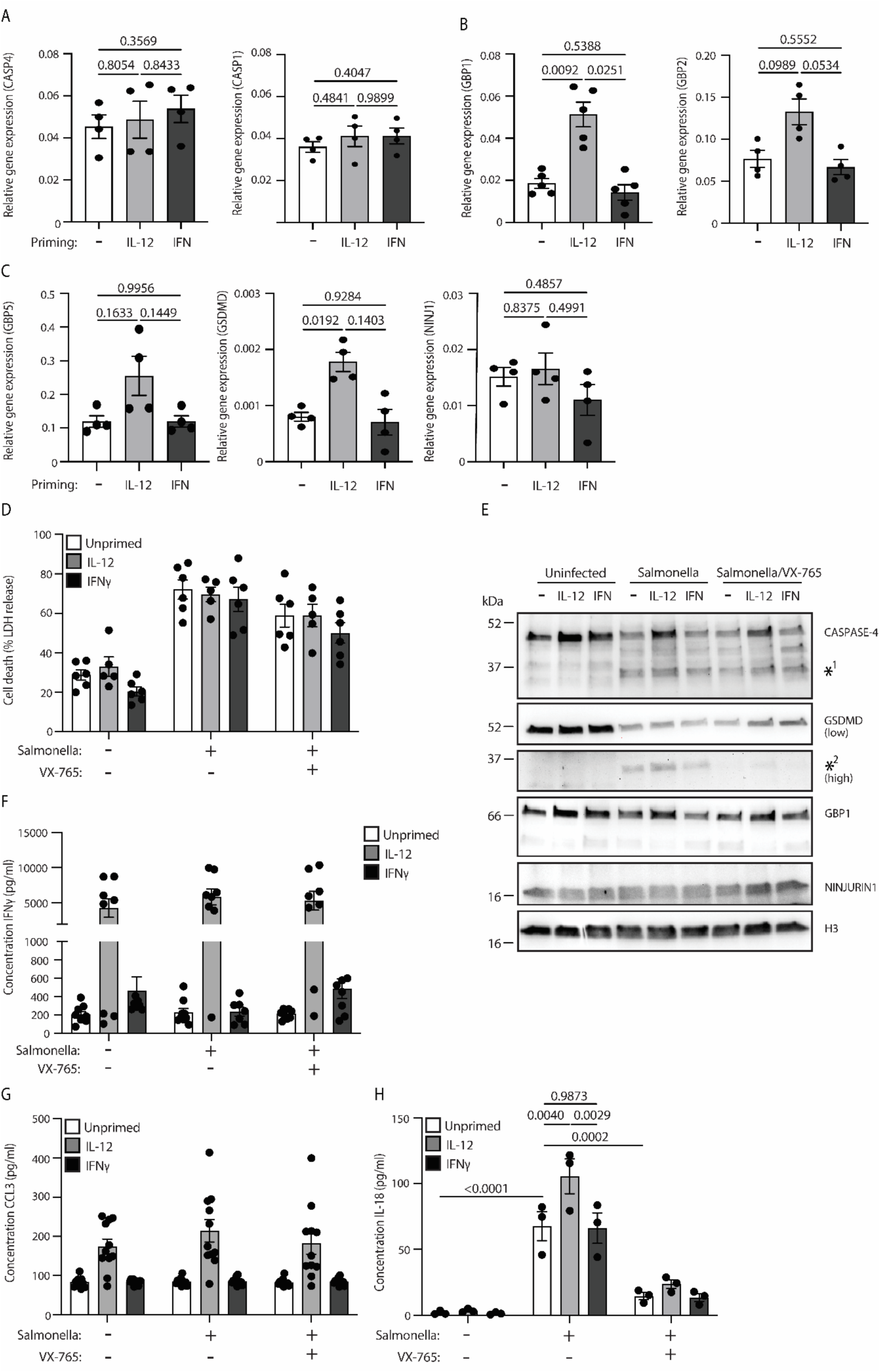
IL-12 but not IFN*γ*-mediated activation of human blood NK cells leads to upregulation of non-canonical inflammasome components. Expanded human peripheral blood NK cells were treated with and without rhIL-12 or rhIFN*γ* for 16 hrs. Quantitative PCR was used to assess expression of (**A**) *CASP4* and *CASP1*, (**B**) *GBP1* and *GBP2*, and (**C**) *GBP5*, *GSDMD*, and *NINJ1*. Post-treatment, NK cells were additionally infected with *S.* Typhimurium alone, or in combination with VX-765. Supernatant was taken at 5 hrs post-infection in order to assess (**D**) cell death by LDH release and (**F**) IFN*γ*, (**G**) CCL3, and (**H**) IL-18 production by ELISA. (**E**) Representative western blot analysis of expanded and primed human NK cells infected with either *S.* Typhimurium alone or in combination with VX-765 for 5 hrs compared to loading controls of H3. *^1^ indicates a p32 cleavage product of caspase-4. *^2^ indicates a p30 cleavage product of GSDMD. Data from 2 - 3 independent experiments, n = 3 - 8 donors. Data in (A, B C and H) were analysed by one-way ANOVA with Dunnett’s multiple-comparison test. p values are displayed and p<0.05 was considered statistically significant.

## Discussion

Inflammasome pathways have been extensively characterised in macrophages and epithelial cells, yet whether lymphocytes engage these signalling platforms remains unclear. Here, we demonstrate that human NK cells possess a functional non-canonical inflammasome (NCI) that enables cell-intrinsic detection of cytosolic Gram-negative bacteria. This finding repositions NK cells from passive recipients of inflammasome-derived cytokines to autonomous and active participants in anti-bacterial immunity. The NK cell NCI displays a species-specific protein profile and is primed by IL-12 rather than IFNγ.

Human NK cells use a limited inflammasome system. They lack the canonical sensors NLRP3 and NLRC4, do not express TLR4, and fail to produce IL-1β upon bacterial infection. Yet they retain functional caspase-4, guanylate-binding proteins (GBP1, GBP2, GBP5), and GSDMD, enabling detection of cytosolic LPS and execution of pyroptosis. The NK NCI produces a specific output profile with IL-18 secretion and inflammatory cell death without IL-1β production, highlighting fundamental differences from macrophage responses to the same pathogen. However, the prevalence of such mechanisms across lymphocyte populations remains to be fully investigated. Previous work suggested that T cell subsets activate the canonical CARD8 inflammasome upon treatment with chemotherapeutics, triggering cell death without cytokine production (38).

Strikingly, murine NK cells, despite expressing inflammasome component transcripts at levels comparable to macrophages, lack functional caspase-11 protein and do not undergo pyroptosis upon *Salmonella* infection. This transcript-protein discordance suggests post-transcriptional or translational regulation that differs between species. This species difference has practical implications: mouse models have been used extensively to study NK cell responses to *Salmonella*, including in cancer immunotherapy contexts (64). Our findings urge caution in extrapolating from these studies to human NK cell biology and suggest that human-specific experimental systems are essential for understanding the contributions of inflammasomes to human NK cell function.

The selective responsiveness of human NK cells to IL-12 rather than IFNγ priming represents an unexpected regulatory distinction from other cell types. IFNγ, signalling through JAK-STAT1 (65), is the canonical priming agent for non-canonical inflammasome components in macrophages and epithelial cells (9–11). In NK cells, however, IFNγ had no effect on caspase-4, GBPs, or GSDMD expression. Recent transcriptomic analyses support this interpretation, demonstrating that NK cells show minimal response to IFNγ compared with T and B cells (66). IL-12, via STAT4 signalling (61), significantly upregulated these components and enhanced infection-induced IL-18 secretion. This suggests that the promoters of key NCI genes may harbour STAT4-responsive elements that are preferentially active in NK cells. The IL-12 dependency also aligns with clinical observations that individuals with defects in IL-12 signalling are particularly susceptible to *Salmonella* infection, raising the possibility that impaired NK cell inflammasome priming contributes to this phenotype (60,63).

Our observation that *Salmonella* infection induces IL-18 secretion from NK cells indicates a potential for autocrine amplification of NK cell responses. IL-18 receptor expression varies across NK cell subsets, with recent single-cell analyses indicating the highest expression in NK2 (CD56high, CD16low) populations. If NCI-competent NK cells (predominantly NK1 and NK3, based on our expression data) secrete IL-18 upon infection, this could preferentially activate nearby NK2 cells, thereby amplifying the local immune response across tissues. Such subset-specific interactions deserve investigation, as they may shape NK cell responses to bacterial infection. More broadly, the heterogeneity of inflammasome component expression across NK cell subsets with PYCARD/ASC expression being restricted to NK1B and NK1C, highlighting that different NK populations may have distinct capacities for inflammasome activation, a complexity that functional studies have yet to address.

Our finding that a subset of NK cells remains infected with *Salmonella* over 14 days raises questions about bacterial persistence and potential immune evasion. *Salmonella* persistence in macrophage vacuoles is well documented and enables antibiotic evasion and cellular reprogramming (67). Whether similar mechanisms operate in NK cells and whether persistent infection affects NK cell function or memory-like phenotypes remain to be determined. Our data indicate that, at least during the initial infection, *Salmonella* can invade NK cell cytosol and activate the NCI. The absence of NLRP3 and NLRC4 in these cells suggests that cytosolic LPS is likely the primary PAMP sensed by NK cells upon *Salmonella* infection. However, we cannot exclude other inflammatory conditions that may prime caspase-1 activation in these cells (e.g., other pathogens).

In conclusion, we have identified a specialised non-canonical inflammasome pathway in human NK cells that operates independently of canonical inflammasome components and is regulated by IL-12 rather than IFNγ. This pathway enables NK cells to directly sense intracellular Gram-negative bacteria, adding to the network of innate immune cells capable of autonomous pathogen detection. The species-specific nature of this response highlights the importance of human experimental systems for understanding NK cell inflammasome biology. Future work should address how this pathway shapes tissue responses to infection and whether it contributes to diseases such as sepsis, in which reduced NK cell numbers are observed and are associated with poor patient outcomes (68).

## Material and Methods

### Study approval

Work with human primary cells was performed under approval by the University of York Biology Ethic Committee (DB202111). Mice were bred and maintained in specific pathogen-free conditions in the University of York, Biological Services Facility. Surplus C57 BL/6 mouse tissue was used with the approval of the local AWERB committee, and under license with the UK Home Office.

### Human immune cell isolation and expansion

Human NK cells, PBMCs and neutrophils were isolated from fresh blood in EDTA-coated tubes (BD Bioscience) or from blood cones (apheresis product, NHS) collected from anonymous healthy donors. Neutrophils (only from fresh blood) and NK cells were isolated from whole blood by negative selection using the EasySep Direct Neutrophil isolation kit (StemCell Technology, 19666) and EasySep Direct human NK isolation kit (StemCell Technology, 19665). To ensure a maximal NK purity, isolated NK cells were expanded for 14 days with Immunocult NK Cell Expansion Kit (StemCell Technology, 100-0711), according to manufacturer instructions. After 14 days, dead cells from NK culture were removed using EasySep Dead Cell Removal (Annexin V) Kit.

PBMCs were isolated from whole blood using Lymphocyte Separation Medium (Lonza, BESP1109E). Red blood cells were lysed using ACK lysis buffer (Gibco, A1049201) for 5 min at room temperature. PBMCs were differentiated into macrophages with 50 ng/ml rhM-CSF (Proteintech, HZ-1192) in RPMI, 10% FBS, 1% penicillin/streptomycin, and 1% glutamine for 7 days at 37°C, 5% CO_2_.

### Murine immune cell isolation and priming

C57BL/6J mouse spleens were collected in PBS with 2% FBS. Single cell suspensions were obtained by passing tissue through a 70µm cell strainer (Corning, 352350). Red blood cells were lysed using ammonium chloride solution for 5 minutes at room temperature. Neutrophils were isolated using an EasySep Direct Neutrophil isolation kit (StemCell Technology, 19855). NK cells (LiveCD45^+^CD3CD4CD8CD19CD11bCD11cFcεR1Gr1 (lineage)^-^NK1.1^+^) and macrophages (LiveCD45^+^CD3^-^CD11b^high^) were stained with flow cytometry antibodies (see **Supplementary Table 1**) and isolated by FACS using a MoFlo Astrios (Beckman Coulter, Summit software). Mouse NK cells and macrophages were primed with 100 ng/mL of LPS EK (Invivogen) for 3hrs or used directly for RNA extraction after isolation.

### Bacterial infection, cell priming, and cell death analysis

NK cells, neutrophils, and differentiated macrophages were plated in 12-well plates in Opti-MEM (GIBCO) at 1×10^6^ cells/ml, 2.5×10^6^ cells/m and 0.5×10^6^ cells/ml, respectively. Cells were infected with either *Salmonella enterica* Serovar Typhimurium SL1344 or LT2, or *S. entericaΔorgA*, LT2-TdTomato, LT2-pPuhpT-GFP, or *Staphylococcus aureus* grown in log phase. Briefly, bacterial strains were cultured in lysogeny broth (LB) medium overnight at 37°C on an orbital shaker. These were subcultured 1:40 and grown for 3h at 37°C to generate log phase cultures. Before infection, bacteria were collected by centrifugation, and washed twice with pre-warmed PBS. Cells were infected at MOI 50 for 1h at 37°C. After 1 hr, gentamicin was added at a final concentration of 100 µg/ml to kill extracellular bacteria and cells were incubated for 4 hrs at 37°C, 5% CO_2_. As indicated, VX-765 (25 µM in DMSO, Sigma-Aldrich, 5.31372) was added 30 mins before inflammasome activation.

In some experiments, NK, rested for 6 hours in NK medium without NK cell Expansion Supplement (StemCell Technology, 100-0711) and neutrophils were primed 3 hours with 100 ng/ml ultrapure LPS-EK (Invivogen, tlrl-peklps) or with 25 ng/ml rhIFNγ (Proteintech, HZ-1301) or 100 ng/ml rhIL-12 (Proteintech, HZ-1256) for 16 hours. NLRP3 inflammasome was activated using Nigericin (20 µM in EtOH, Sigma-Aldrich, N7143) for 3 hrs at 37°C. As indicated MCC950 (10 µM in H_2_O, Sigma-Aldrich, PZ0280) were added to the cell culture media 30 min before inflammasome activation. NLRP1 was activated using 3 µM Val-boroPro for 6 hrs at 37°C (Vbp, 5.31465).

For cell death measurements lactate dehydrogenase release in the supernatants was measured using a CytoTox 96® Non-Radioactive Cytotoxicity Assay (Promega, G1780). Absorbance was measured on a plate reader (Infinite M Plex, TECAN) at 490nm with a 650nm background subtraction following manufacturer’s instructions.

### Quantitative PCR analysis

mRNA from primary cells (mouse and human) were isolated using the Arcturus PicoPure Isolation kit (ThermoFisher, KIT0204) and cDNA was generated using the SuperScript IV First Strand kit (ThermoFischer, 18091050), using manufacturer’s instructions. qPCR was performed using TaqMan probes (Life Technologies, **Supplementary Table 1**) on an AriaMx Real-time PCR system (Agilent).

### Immunoblotting

Cell extracts were lysed with cell sample buffer (2% SDS, 66mM Tris pH7.6 reconstituted in NuPage 1X LDS buffer (Novex, P5878547) and run on 10% or 4 - 20% acrylamide gels (BioRad) with methanol-chloroform precipitated cell supernatant mixes (69). Gels were transferred on 45 µM nitrocellulose membrane using the TurboBlot transfer system (BioRad), and blocked in 5% skimmed milk TBST. Samples were then incubated overnight at room temperature in the primary antibody at 4°C and for 2 hrs the following day in the secondary antibody (**Supplementary Table 1**). Membrane was imaged using the Immobilon Forte ECL Substrate (VWR, WBLUF) using a ThermoFisher iBright 1500.

### Flow cytometry

Single cell suspensions were incubated with anti-CD16/CD32 antibodies (Biolegend, Clone 93) followed by fluorochrome-conjugated antibodies as detailed in **Supplementary Table 1**. All samples were co-stained with a cell viability dye (Zombie YG581 or Aqua, Biolegend, 423123 and 423102). In some experiments with human blood NK cells after *Salmonella* infection, and following the addition of gentamicin, YVAD-FMK FLICA660 or FLICA-FAM was added following manufacturer instructions (1:1000, FLICA Caspase-1 kit, BIO-RAD, 9120 and ICT098) and cells were incubated for 1 hr at 37°C. Cells were washed twice with the 1x kit wash buffer. Untreated NK cells stained for FLICA were used as control. Sample acquisition used an LSR Fortessa, CytoFLEX LX (Beckman Coulter), or FACSDiscover™ S8 (BD Biosciences, CellView™ Image) and then visualised using FlowJo (version 10.10) software. For cell sorting, an Astrios (70 µm nozzle, Beckman Coulter, Summit software) was used.

### Immunofluorescence microscopy

8-well chamber slides (Ibidi, 80821) were coated with 12.5 µg/ml fibronectin for 1 hr, washed with PBS and transferred to 4°C until use. NK cells were seeded directly into chamber slides at 4×10^5^ cells/well and immediately infected with log phase *Salmonella* LT2-TdTomato at MOI 50 for 1h at 37°C, 5% CO_2_. Gentamicin was added directly to the culture medium, and cells were cultured for 4 hrs at 37°C, 5% CO_2_. Cells were subsequently fixed with 4% PFA (in PBS) for 10 min at room temperature and washed with PBS. Cells were blocked with 5% goat serum for 2 hrs, stained with rabbit anti-human NCAM (CD56, Cell Signalling, 99746) overnight, washed with PBS and stained with goat anti-rabbit IgG (H+L), AF488 conjugate (Cell Signalling, 4412) for 1 hr. Cells were washed with PBS and stained with DAPI (2µg/ml) (Biotum, 40043), ready for imaging on Zeiss Elyra 7.

### Cytokine assays

Cytokine secretion in cell culture supernatants was assessed using standard sandwich ELISA, IL-1β (Invitrogen, 88-7261), CCL3 (Invitrogen, 88-7035), IL-18 (R&D systems, DY318-05), GM-CSF (Invitrogen, 88-8337), IL-22 (Invitrogen, 88-7522), according to the manufacturer’s instructions. IFNγ secretion was measured using the Lumit IFNγ immunoassay (Promega, W6040) following the manufacturer’s instructions in a 384 well plate. Luminescence was measured on a plate reader (Infinite M Plex, TECAN).

### Bioinformatical data analysis and statistics

Mouse RNA sequencing data was obtained from the GEO database (GSE143943, (49). The data was filtered for genes relevant to NCI and the mean expression values log transformed. Heat maps were generated in R Studio using the pheatmap package. Clustering was automatically applied to the gene list.

Human scRNAseq datasets were obtained from either the foetal immune cell atlas (https://developmental.cellatlas.io/#) or peripheral blood (https://storage.googleapis.com/haniffalab/meta-nk/PBMC_Noreg.h5ad) and mined to analyse tissue resident or blood NK cells, respectively. The published data was loaded with *scanpy* and the inflammasome component genes visualised in a violin or *dotplot*. The dotplot was grouped by the 6 NK clusters discovered in the original study, and the genes were standard-scaled across these clusters to visualize relative abundance of each gene across cell contexts.

Statistical analysis for most experiments was performed using Prism10 (GraphPad, San Diego, CA).

## Acknowledgement

D.B. is supported by an MRC New Investigator Award (MR/Z504221/1), a Springboard Award (Academy of Medical Sciences and Wellcome Trust) and a Royal Society Grant. J.A. was supported by a GenerationResearch MSc Studentship and VISFO. J.L.B., D.G.K, and M.C. are supported by the NIHR Leeds Biomedical Research Centre (BRC) (NIHR203331), and J.L.B and D.G.K. by an MRC National Mouse Genetics Network grant (MC_PC_21043). D.G.K is additionally supported by a CRUK Programme Foundation Award (DCRPGF\100008), an MRC-AMED joint award (MR/V005502/1), and Blood Cancer UK (24014). M.G. was in receipt of a BBSRC White Rose DTP PhD Studentship (BB/T007222/1).

W.G. is supported by an MRC Career Development Fellowship (MR/X007146/1), Leukaemia U.K. (2021/JGF/002), an EHA bilateral collaborative award and the Children’s Cancer and Leukaemia Group (CCLGA 2022 13 Grey). K.S.R is supported by the Academy of Medical Sciences Springboard (SBF009\1153), Royal Society (RG\R1\241058) and Hull York Medical School Start-up. The authors thank the Centre for Blood Research (CBR), SUKIT and the InflammaZoom UK networks for helpful discussions and constructive criticism on this research. The authors thank the phlebotomist team at the University of York for their support with collecting fresh blood for experimental purposes and the donors who kindly provided the blood. N/TERT was a kind donation from Prof. James Rheinwald (70). We would like to thank Prof. Petr Broz (University of Lausanne) and Prof. Marjan van der Woude (University of York) for providing the bacterial strains used in this study.

## Authors contribution

N.G.F.L, J.A, A.H, Z.A, K.S.R, M.G, C.R, A.V, J.L.B, M.A.C, F.P. and D.B performed experiments. N.G.F.L, Z.A, J.H, J.L.B and D.B designed experiments, D.B secured funding, supervised, designed and oversaw the project. N.G.F.L, J.A, A.H, K.S.R, K.H, G.C, J.L.B, D.B analysed data. D.B, N.G.F.L, and J.L.B wrote the paper with inputs from all authors.

## Supplementary Figure Legends

**Supplementary Figure 1.**
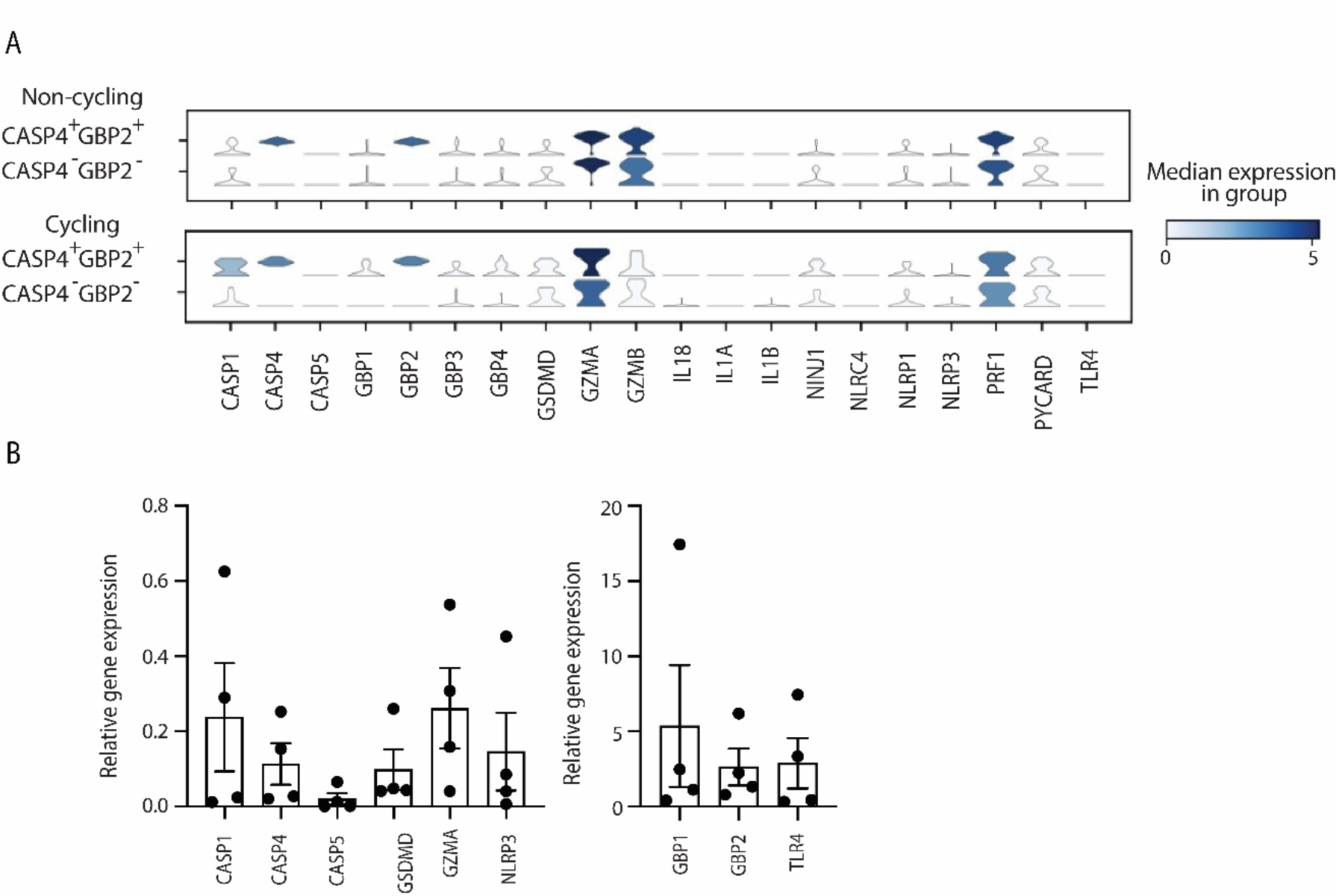
Human thymic NK cells express components of the non-canonical inflammasome. (**A**) scRNA-seq data of inflammasome components in *CASP4^+^GBP2^+^* and *CASP4^-^GBP2^-^* human foetal thymic non-cycling and cycling NK cells showing the median expression and number of cells expressing inflammasome components. (**B**) Quantitative PCR analysis of non-canonical inflammasome components in neutrophils, n = 4 donors, matched with NK cell phenotypes in Fig. 1B.

**Supplementary Figure 2.**
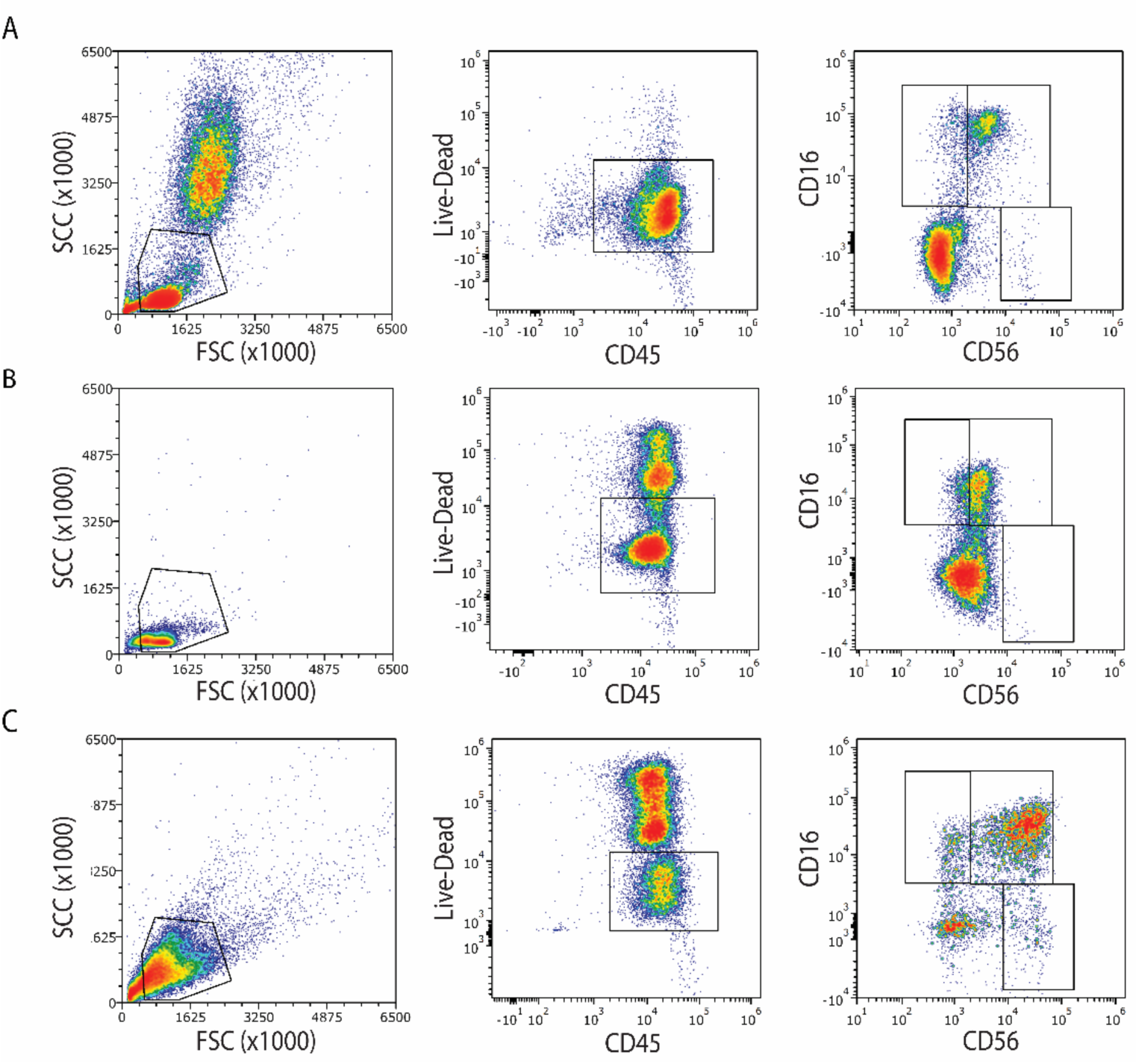
Characterisation of NK cell subpopulations in human blood. Representative density plots of (**A**) PBMCs, (**B**) negatively-selected NK cells and (**C**) 14 days expanded NK cells from human peripheral blood showing typical percentages of CD56^int^CD16^+^ and CD56^high^CD16^-^. Human peripheral blood from (**C**) before and (**D**) after NK cell negative selection, analysed by flow cytometry to show typical percentages of CD56^int^CD16^+^ and CD56^high^CD16^-^, and within this NK1, NK2, and NK3 subsets, as defined by Rebuffet *et al* (2024).

**Supplementary Figure 3.**
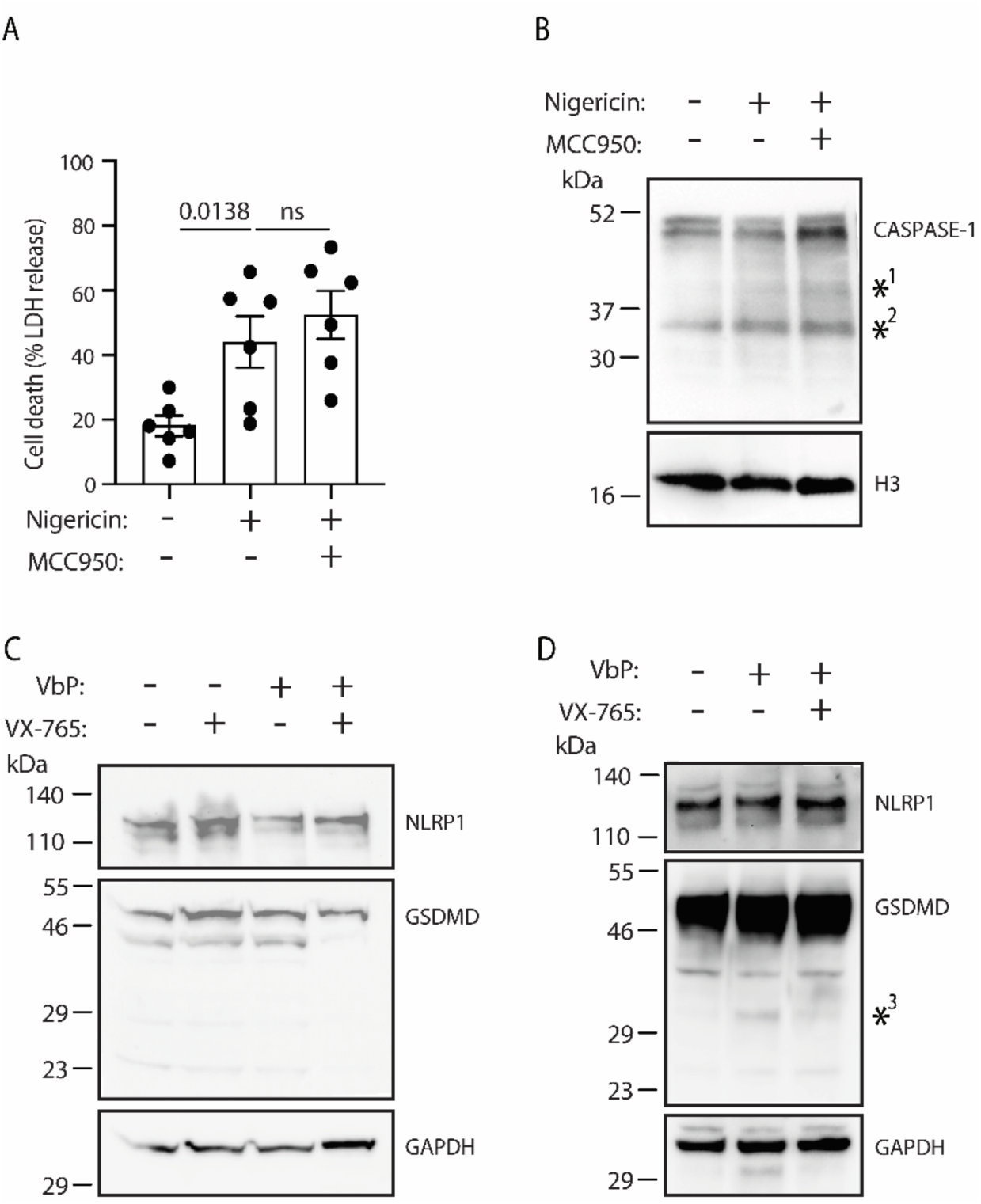
NLRP1, NLRP3, and FLICA activation in human blood NK cells. Expanded human peripheral blood NK cells were treated with nigericin (20 µM), with or without MCC950 (10 µM) for 3 hrs. Cells were then assessed for (**A**) the release of LDH for cell death and (**B**) the cleavage of caspase-1 by western blot. (**C**) NK cells or (**D**) N/TERT keratinocytes were treated with Val-boroPro (3µM, VbP) for 6hrs, with and without VX-765, and assessed by western blot. n = 4 - 6 donors. *^1^ and *^2^ indicate possible NK cell-specific protein isoforms of CASPASE-1. *^3^ indicates the p30 cleavage fragment of GSDMD. Data in (A) were analysed by one-way ANOVA with Dunnett’s multiple-comparison test. p values are displayed and p<0.05 was considered statistically significant.

**Supplementary Figure 4.**
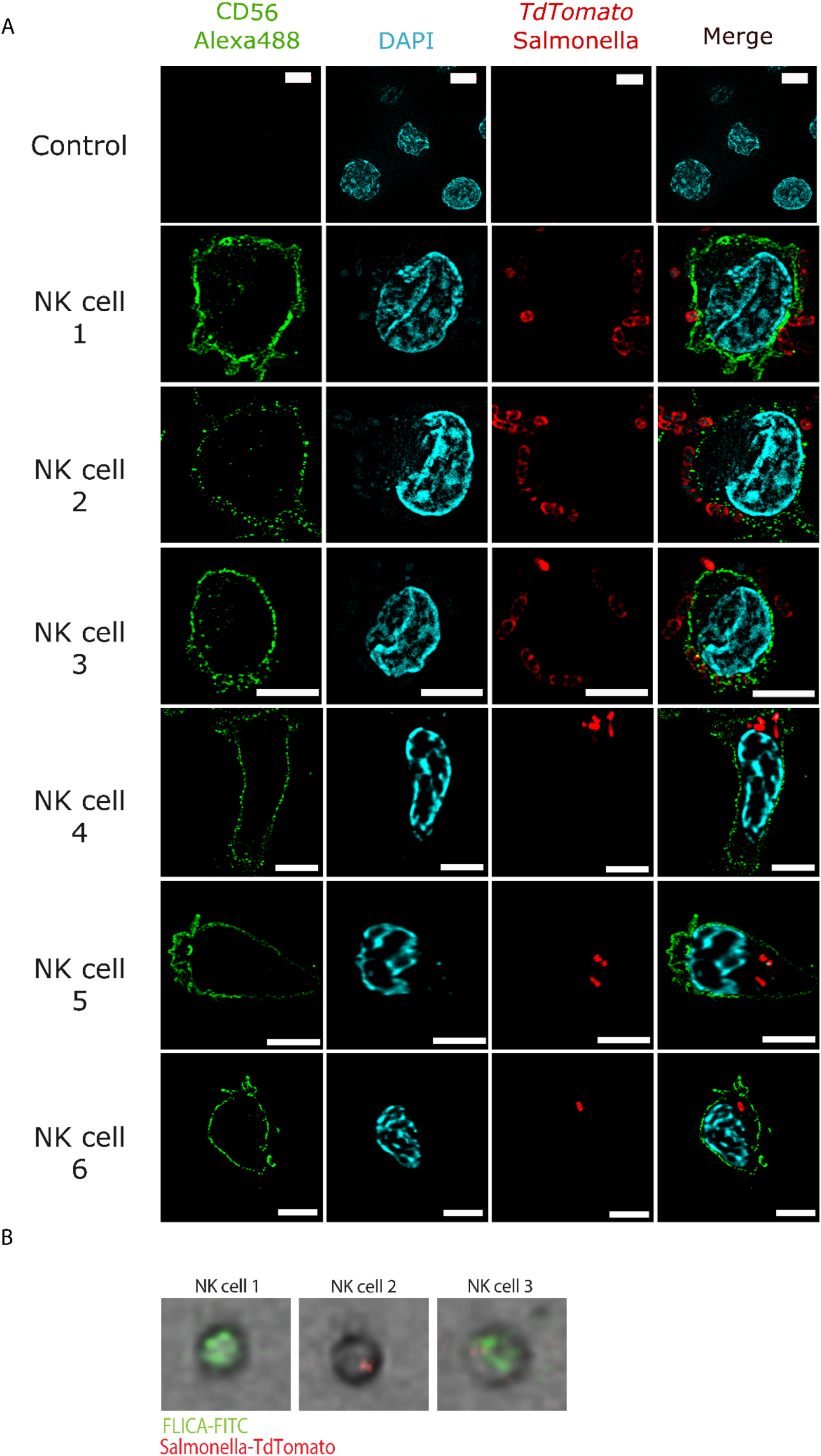
*Salmonella* Typhimurium interacts with and enters NK cells, activating inflammatory caspases. Expanded human peripheral blood NK cells were infected with *S.* Typhimurium-TdTomato. (**A**) At 5 hrs post-infection NK cells were stained on fibronectin coated slides with anti-CD56 and DAPI and visualised by fluorescence microscopy (white bar indicates 5 µm). Panel shows range of NK cell-*Salmonella* interactions. Control represents non-infected NK cells stained with anti-CD56 and DAPI. (**B**) Representative images of CD56^+^ cells were reconstructed, using CellView^TM^ beam-splitting technology, to show human NK cells that were FLICA-FAM^+^*Salmonella-* TdTomato^-^ (NK cell 1), FLICA-FAM^-^*Salmonella-*TdTomato^+^ (NK cell 2), and FLICA-FAM^+^*Salmonella-*TdTomato^+^ (NK cell 3), respectively.

**Supplementary Figure 5.**
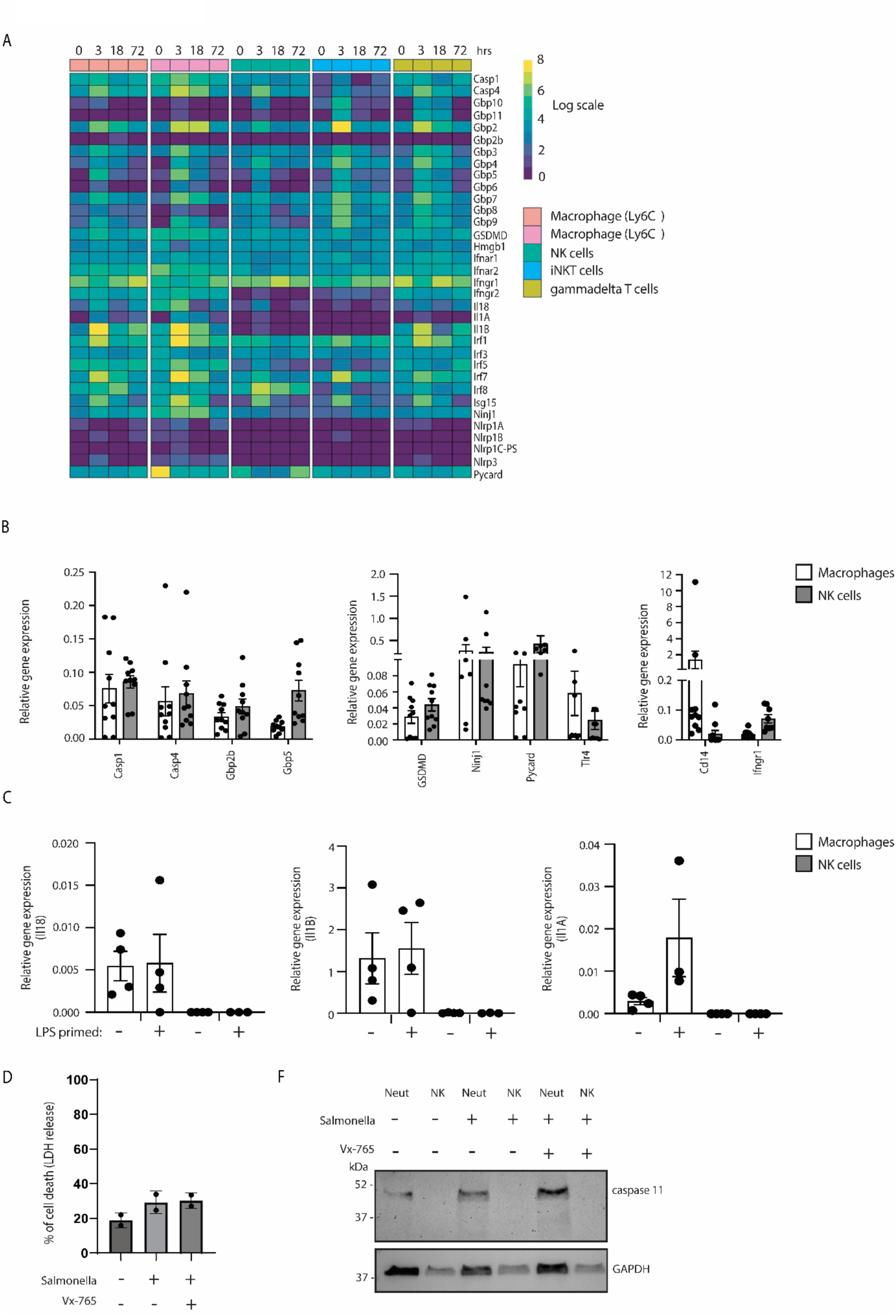
Canonical and non-canonical inflammasome components are expressed in mouse NK cells at the gene, but not the protein level. (**A**) RNASeq dataset extracted from flow cytometry sorted immune cells showing a heatmap expression analysis of canonical and non-canonical inflammasome-associated genes (58). Murine spleen NK cells and CD11b^high^ cells were sorted and cDNA was extracted from either (**B**) naive populations or (**C**) from LPS (100 ng/mL, 3 hrs)-primed cells. Expression of canonical and non-canonical inflammasome-associated genes was assessed by quantitative PCR using *Gapdh* as a house-keeping gene, n = 4-8, collected across 2 independent experiments. (**D**) Representative western blot analysis of caspase-11 expression in LPS-primed murine spleen NK cells.

**Supplementary Table 1.** (A) Quantitative PCR Taqman primer sets for human and mouse targets. (B) Antibodies used in flow cytometry (FC), immunofluorescence (IF), and western blot (WB).

| Human gene | Taqman ID | Mouse gene | Taqman ID |
| --- | --- | --- | --- |
| *NLRP3* | Hs00918062 | *Ninj1* | Mm00479014 |
| *NLRC4* | Hs00892666 | *Gsdmd* | Mm00509958 |
| *NINJ1* | Hs00982607 | *Il1a* | Mm00439620 |
| *TLR4* | Hs00152939 | *Il1b* | Mm00434228 |
| *GSDMD* | Hs00226875 | *Il118* | Mm00434225 |
| *Casp1* | Hs003354836 | *Gbp2b* | Mm00657086 |
| *Casp4* | Hs01031951 | *Gbp5* | Mm00463735 |
| *Casp5* | Hs07290189 | *Ifngr1* | Mm00599890 |
| *GBP1* | Hs00266717 | *Cd14* | Mm00438094 |
| *GBP2* | Hs00894837 | *Tlr4* | Mm00445273 |
| *GBP5* | Hs00369472 | *Casp1* | Mm00438023 |
| *GzmA* | Hs009869184 | *Casp4* | Mm00432304 |
| *IFNGR1* | Hs00166223 | *Pycard* | Mm00445747 |
| *GAPDH* | Hs02758991 |  |  |
| *IL1B* | Hs01555410 |  |  |

**B**
| Human antibody target | Clone number | Catalog number | Supplier | Use |
| --- | --- | --- | --- | --- |
| CASPASE-1 | D7F10 | 3866 | Cell Signalling | WB |
| GBP1 | polyclonal | 15303-1-AP | Proteintech | WB |
| GBP2 | polyclonal | 11854-1-AP | Proteintech | WB |
| GBP5 | polyclonal | 13220-1-AP | Proteintech | WB |
| αTubulin | 1E4C11 | 66031-1-Ig | Proteintech | WB |
| CASPASE-4 | 1D1E12 | 67398-1-Ig | Proteintech | WB |
| GAPDH | 1E6D9 | 60004-1-Ig | Proteintech | WB |
| NINJURIN 1 | 2G5 | H00004814-MO1A | Abnova | WB |
| GSDMD | E9S1X | 39754S | Cell Signalling | WB |
| NLRP1 | 9F9B12 | 679802 | Biologend | WB |
| H3 | 1A2A3 | 68345-1-Ig | Proteintech | WB |
| CD45 BUV395 | HI30 | 363-0459-42 | Thermo | FC |
| CD4 PE | SK3 | 60122PE | StemCell Technologies | FC |
| CD8a PE | HIT8a | 300908 | Biologend | FC |
| CD19 PE | HIB19 | 302208 | Biologend | FC |
| CD14 PE | HCD14 | 325605 | Biologend | FC |
| CD123 PE | 6H6 | 306006 | Biologend | FC |
| CD303 PE | 201A | 354204 | Biologend | FC |
| CD3e BV605 | UCHT1 | 300459 | Biologend | FC |
| CD16 BV480 | eBioCB16 | 414-0168-41 | Thermo | FC |
| CD56 BUV661 | TULY56 | 376-0566-42 | Thermo | FC |
| CD127 PECy7 | R34.34 | 25-1271-82 | Thermo | FC |
| CD2 AlexaFluor700 | RPA-2.10 | 300238 | Biologend | FC |
| CD57 BV421 | QA17.A04 | 393326 | Biologend | FC |
| NCAM1 (CD56) | E7X9M | 99746 | Cell Signalling | IF |
| Mouse antibody target | Clone Number | Catalog Number | Supplier | Use |
| Caspase-11 | 17D9 | NB120-10454 | Novus Biologicals | WB |
| GBP2 | polyclonal | 11854-1-AP | Proteintech | WB |
| H3 | 1A2A3 | 68345-1-Ig | Proteintech | WB |
| CD45 AlexaFluor700 | 30-F11 | 103128 | Biologend | FC |
| CD3 PE | 17A2 | 100206 | Biologend | FC |
| CD4 PE-Cy7 | GK1.5 | 100421 | Biologend | FC |
| CD8a PE-Cy7 | 53-6.7 | 100721 | BioLegend | FC |
| CD19 PE-Cy7 | 1D3 | 25-0193-82 | eBioscience | FC |
| CD11b BV510 | M1/70 | 101263 | Biologend | FC |
| CD11c PE-Cy7 | N418 | 117317 | BioLegend | FC |
| FcεR1a PE-Cy7 | MAR-1 | 134317 | BioLegend | FC |
| Gr1 PE-Cy7 | RB6-8C5 | 108416 | BioLegend | FC |
| NK1.1 BV421 | PK136 | 108732 | Biologend | FC |
| Other antibodies | Clone Number | Catalog Number | Supplier | Use |
| Mouse Ig-HRP | Polyclonal | RGAM001 | Proteintech | WB |
| Rabbit Ig-HRP | Polyclonal | RGAR001 | Proteintech | WB |
| Anti-rabbit IgG (H+L),<br>(Alexa Fluor® 488 Conjugate) | F(ab')<sub>2</sub> Fragment | 4412 | Cell Signalling | IF |

## Notes

### Competing Interest Statement

The authors have declared no competing interest.

### Summary of Updates

Additional results have been added regarding the murine NK cells to further support our conclusions. We have added additional microscopy images as well of infected NK cells.

